# PKMζ in the nucleus accumbens acts to dampen cocaine seeking

**DOI:** 10.1101/320945

**Authors:** Anna G. McGrath, Jeffrey D. Lenz, Lisa A. Briand

**Author notes:** **Address correspondence to:** Lisa A. Briand, Ph.D., Temple University Department of Psychology, Weiss Hall, 1701 North 13^th^ Street Philadelphia, PA 19122.

## Abstract

The constitutively active, atypical protein kinase C, protein kinase M-ζ (PKMζ), is exclusively expressed in the brain and its expression increases following exposure to drugs of abuse. However, the limitations of currently available tools have made it difficult to examine the role of PKMζ in cocaine addiction. The current study demonstrates that constitutive deletion of PKMζ potentiates cue-induced reinstatement of cocaine seeking and increases both food and cocaine taking, without affecting cue-driven food seeking in both male and female mice. Conditional deletion of PKMζ within the nucleus accumbens recapitulated the increase in cocaine taking and seeking seen in the constitutive knockout mice, but only in male animals. Site-specific knockdown of PKMζ in the nucleus accumbens had no effect on cocaine or natural reward behaviors in female mice. Taken together these results indicate that PKMζ may act to dampen addictive phenotypes. Furthermore, these results indicate that PKMζ is playing divergent roles in reward seeking in males and females.

## Introduction

PKMζ, a constitutively active form of protein kinase C, has been extensively studied for its supposed role in memory maintenance. Early work in this area demonstrated that application of PKMζ is sufficient to induce long-term potentiation (LTP) *in vitro* (Ling *et al*, 2002). Stimulation protocols to induce LTP also increase PKMζ (Hrabetova and Sacktor, 1996; Sacktor *et al*, 1993). Furthermore, PKMζ has been shown to be involved in memory processes *in vivo*. PKMζ is upregulated following learning and memory tasks (Hsieh *et al*, 2017; Li *et al*, 2011). Inhibition of PKMζ has been shown to disrupt memory maintenance. Administration of a PKMζ inhibitor, Zeta Inhibitory Peptide (ZIP), into the hippocampus blocked the ability of rats to recall a previously learned fear response 24hrs later (Pastalkova *et al*, 2006). Similar, long-lasting disruption of memories has been seen with spatial memory following ZIP infusion into the hippocampus (Pastalkova *et al*, 2006; Serrano *et al*, 2008), taste memory following ZIP infusion in the insular cortex (Shema *et al*, 2007), fear memory following ZIP infusions in the amygdala (Migues *et al*, 2010; Serrano *et al*, 2008) and conditioned reward following ZIP infusions into the nucleus accumbens (Li *et al*, 2011).

More recent data has shed doubt on the necessity of PKMζ for learning and memory. No LTP or memory deficits were found in two independently generated lines of PKMζ knockout mice (Lee *et al*, 2013a; Volk *et al*, 2013). Furthermore, when given ZIP, PKMζ knockout mice show memory impairments (Lee *et al*, 2013a). These findings indicate that the memory dampening effects of ZIP are not due to its actions on PKMζ. However, the ability of experience to upregulate PKMζ expression suggests it may still be playing a role in behavioral and synaptic plasticity. Since many past experiments utilized the non-specific inhibitor, ZIP, previous studies examining the role of PKMζ are difficult to interpret.

Despite the mixed evidence for the importance of PKMζ in memory maintenance, there does appear to be a role for PKMζ in motivation and reward processing. PKMζ KO mice consume more alcohol than their wildtype controls in an intermittent access procedure (Lee *et al*, 2014). Very few studies have examined reward-related behaviors in PKMζ KO mice. Therefore, the aim of these studies was to determine if the ability of PKMζ to dampen reward extends beyond the published effects on alcohol. To achieve this we utilized constitutive PKMζ knockout mice to examine the role of PKMζ in cocaine self-administration, extinction and reinstatement of cocaine seeking. Additional studies examined how conditional knockdown of PKMζ within the nucleus accumbens affected cocaine taking and seeking. Our results suggest that PKMζ reduces cocaine taking and seeking and the mechanism of action may be sex-specific.

## Materials and Methods

### Subjects

<u>Wildtype studies</u>: Male and female C57BL/6J mice, bred in house, were utilized for the PKMζ abundance studies. <u>Constitutive deletion</u>: The current study utilized PKMζ knock-out mice as described previously (Volk *et al*, 2013). Heterozygous PKMζ KO mice on a C57BL/6J background were mated resulting in mutant and wildtype littermates. <u>Conditional deletion</u>: Mice homozygous for the Cre/lox-conditional allele of PKMζ (flox/flox) were bred on a C57BL/6J background as described previously (Volk *et al*., 2013). Mice (2-6 months old, 20-40g; age matched across group) were group housed until the start of the behavioral experiments at which point they were individually housed. All animals were housed in a temperature and humidity controlled animal care facility with a 12 hr light/dark cycle (lights on a 7:00 A.M.). All procedures were approved by the Temple University Animal Care and Use Committee.

### Adeno-associated virus constructs

The adeno-associated virus (AAV) expressing Cre recombinase (AAV2/9.CMV.PI.CRE, titer 2.84*10^13^ gc/μl) and the AAV expressing green fluorescent protein (eGFP) (AAV2/9.CMV.eGFP, titer 3.74*10^13^ gc/μl) were generated by the Univ. of Pennsylvania Vector Core. AAVs were diluted in sterile phosphate-buffered saline (PBS) for microinjections.

### Intraaccumbal microinjections

PKMζ_flox/flox_ mice (6-8 weeks) were anesthetized with isoflurane and AAV (0.5µL, 1*109 GC/μl) was injected into the accumbens through a 30 gauge needle at a rate of 0.1 µl/min. Stereotaxic coordinates for the nucleus accumbens are (from Bregma) anterior-posterior 1.5, lateral +/− 1.0, dorso-ventral −5.0. Following recovery, mice remained in the home cage for 6 weeks prior to behavioral testing. Viral targeting was confirmed via western blot and mice with PKMζ levels higher than 75% of control levels were removed from the study (n=1).

### Drugs

Cocaine was obtained from the National Institutes of Drug Abuse Drug Supply Program (Bethesda, MD) and dissolved in sterile 0.9% saline.

### Operant Food Training

Prior to catheterization, mice were trained to perform an operant response for sucrose pellets. The mice were placed in operant chambers (Med-Associates) where they learned to spin a wheel manipulandum to receive the sucrose pellet. When the pellet was delivered, a cue light above the active wheel was illuminated, a 2900 Hz tone played, and the house light turned off. This was followed by an 8s time-out where the house light remained off and spinning the wheel had no programed consequences. Mice were able to self-administer up to 50 pellets per 60 min operant session. The mice were food restricted to approximately 90% of their free feeding weight throughout the course of the operant training. They were returned to *ad libitum* feeding three days into the cocaine self-administration sessions.

### Jugular Catheterization Surgery

Mice were anesthetized with 80 mg/kg ketamine and 12 mg/kg xylazine. An indwelling silastic catheter was placed into the right jugular vein and sutured in place. Then the catheter was threaded subcutaneously over the shoulder blade and was routed to a mesh back mount platform (Strategic Applications, Inc.) that secured the placement. The catheters were flushed daily with 0.1 ml of antibiotic (Timentin, 0.93 mg/ml) dissolved in heparinized saline. The catheters were sealed with plastic obturators when not in use.

### Cocaine Self-administration

Mice were given 3-4 days to recover from surgery before beginning behavioral testing. The cocaine self-administration testing was measured over 2 hr sessions (6 days per week) in the same chamber used for the operant food training. During testing, responding on the active wheel delivered an intravenous cocaine injection (0.6 mg/kg/infusion,) paired with the same cues as the food training. After 10 day of cocaine self-administration, cocaine-seeking was extinguished by replacing the cocaine with 0.9% saline. During extinction, the light and tone cues were not present. Daily 2 hr extinction continued until the mice met the extinction criterion of <25% of their responding during the self-administration (average of the last three days). 24 h after meeting the extinction criterion animals underwent a cue-induced reinstatement session. The light and tone were presented non-contingently for 20 s every 2 min during the first 10 min of the session. For the remaining 110 min, the cues were presented following responses on the active wheel, just as was done during the cocaine self-administration sessions. During reinstatement, the mice received saline infusions following active responses. Saline-yoked mice were placed in identical operant chambers however their responding had no programmed consequences. Instead, they received saline infusions along with cue presentations when their partner mouse received cocaine.

### Western Blot

Whole-cell tissue from naïve viral-injected animals was processed for Western blot as described previously (Briand *et al*, 2014). For all samples, protein concentration was quantified using a Pierce BCA Protein Assay Kit (Thermo Scientific). Equal amounts of protein (30 μg) were loaded and separated in 10% Tris-Glycine gels (Invitrogen) and transferred to nitrocellulose membranes using the i-Blot dry transfer system (Invitrogen). Membranes were blocked with Li-Cor blocking buffer. Membranes were incubated for 48 hours at 4 °C with selective antibodies to: PKMζ (1:500 Millipore Sigma SAB4502380) and GAPDH (1:2000, Cell Signaling). Membranes were then incubated with fluorescent secondary antibodies (1:5000, IR-dye 680 or IR-dye 800), before being imaged on an Odyssey fluorescent scanner (Li-cor Biosciences). To ensure equal loading, GAPDH expression was used as a loading control. Separate blots were used for each protein of interest.

### RNA extraction, cDNA synthesis, and quantitative real-time polymerase chain reaction

Mice were killed by cervical dislocation directly from their home cages 14 days following cocaine self-administration or yoked saline experience for the evaluation of early gene expression changes. We chose to examine the nucleus accumbens due to its clear role in cocaine reward and the hippocampus due to the body of previous work demonstrating increases in PKMζ following learning and memory tasks (Hsieh *et al*, 2017). Brains were rapidly removed, whole nucleus accumbens and hippocampi hand-dissected, and frozen in liquid nitrogen. RNA was extracted from hippocampal tissue using TRIzol/chloroform (Invitrogen) and the RNeasy Mini kit (Quiagen). cDNA was synthesized from RNA using an oligo(dT) primer (Operon) and Superscript II reverse transcriptase (Invitrogen). Quantitative real-time polymerase chain reaction (QPCR) was carried out using the SYBR-green master mix (Applied Biosystems) and 300 nM primers (final concentration) on the Stratagene MX3000 using MXPro QPCR software. Cycling parameters were 95°C for 10 min followed by 40 cycles of 95°C (30 sec) and 60°C (1 min), ending with a melting curve analysis to assess the amplification of a single amplicon. All reactions were performed in triplicate, with the median cycle time used for analysis. TATA-box binding protein (TBP) was used as a housekeeping gene against whose levels all experimental genes were normalized. Primer sequences are as follows:

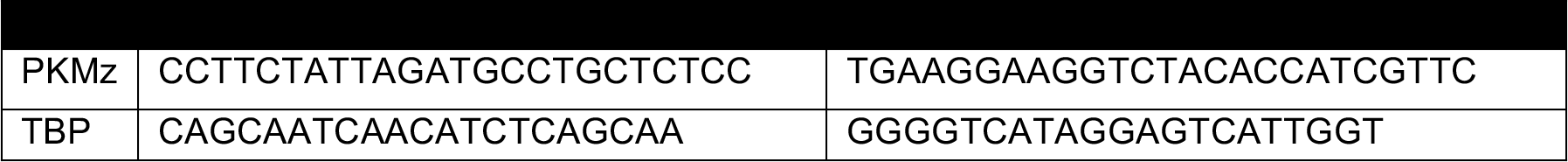

### Data Analysis

All analyses were performed using Graphpad Prism 7.0 software (Graphpad Software). Behavioral data were analyzed using two-tailed Student’s *t*-test, or two-way ANOVA with *Sidak*’s post hoc as appropriate. Statistical significance for all tests was set at α= 0.05.

## Results

### Cocaine Self-Administration and Abstinence Leads to an Increase in PKMζ mRNA and protein abundance within the Nucleus Accumbens

Wildtype mice underwent ten days of training to acquire operant responses for food followed by 10 days of cocaine self-administration (total cocaine intake = 417.6 ± 38.3 infusions). Saline yoked mice were placed in identical operant chambers however their actions had no programmed consequences. They received saline infusions along with cue presentations when their partner received cocaine. Both cocaine-experienced and saline-yoked mice were then placed in their home cages for 14 days prior to brain dissection. Cocaine-self administration experience led to an increase in PKMζ mRNA and protein levels within the nucleus accumbens (mRNA: *t*(8)=5.21, *p*=.0008, n=5-6/group; protein: *t(22)*=2.089, *p*=.048, n=12; Figure 1a,b). No differences were seen in PKMζ mRNA or protein levels in the hippocampus (mRNA: *t*(12)=1.029, *p*=.324, n=5-9/group; protein: *t(22)*=1.50, *p*=.147, n=12-13/group; Figure 1c,d).

**Figure 1.**
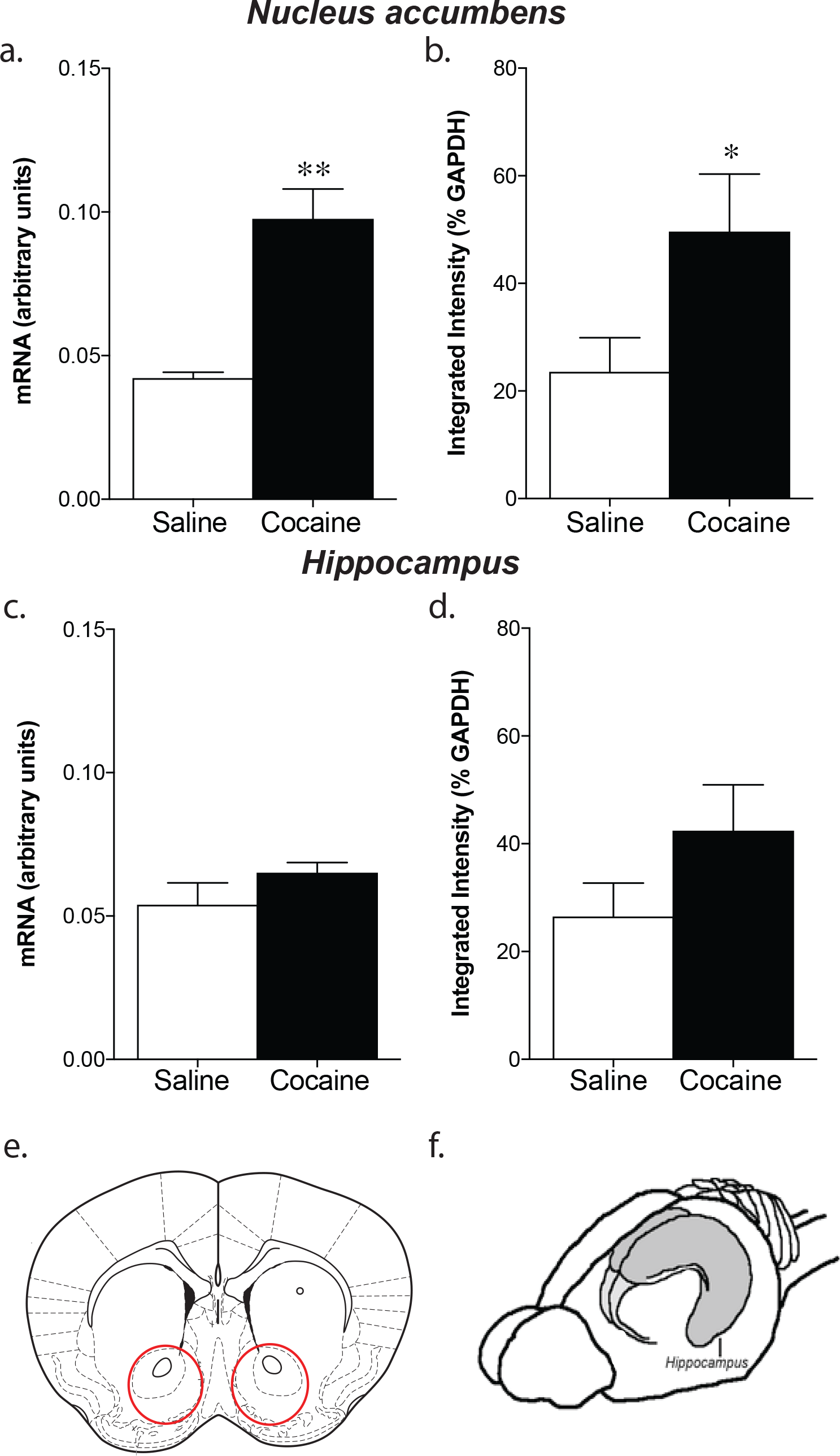
Cocaine self-administration leads to an increase in PKMζ in the nucleus accumbens. After 14 days of abstinence, chronic cocaine self-administration (10 days, 0.5mg/kg/inf, 2hr sessions) leads to an upregulation of PKMζ mRNA (a; *p<.01; n=5-9.group) and protein levels (b; *p<.05; n=12-13/group) within the nucleus accumbens but not the hippocampus (c-d). Red circles indicate area dissected for nucleus accumbens (e). The whole hippocampus (e) was dissected for the measurements of hippocampal mRNA and protein.

### Constitutive Deletion of PKMζ Potentiates Food Self-Administration but does not alter Reinstatement of Food Seeking

Wildtype controls and PKMζ KO_constitutive_ mice underwent ten days of training to acquire operant responses for food. Both wildtype and PKMζ KO_constitutive_ mice showed a gradual increase in the number of pellets earned and active responses per session over the 10 days of food training (effect of session, pellets= *F*(9,342)=7.38, p<.0001; responses= *F*(9, 342)=20.53, p<.0001). Additionally, constitutive deletion of PKMζ, led to an increase in responding for food and increase in the number of rewards received (effect of genotype, pellets= *F*(1,38)=6.79, p=.01; responses= *F*(1,38)=7.15, p=.01; n=22/group; Figure 2a,b). This effect of genotype was seen in both males and females (Males: effect of genotype, pellets= *F*(1,25)=6.54, p=.02; responses= *F*(1,25)=10.15, p=.004; n=10; Females: effect of genotype, pellets= *F*(1,25)=6.75, p=.02; responses= *F*(1,25)=8.04, p=.009; n=10). Both wildtype and PKMζ KO_constitutive_ mice showed an increase in their percent active responding over the 10 sessions but no group differences were seen (effect of session, *F*(9,342)=25.68, p<.0001; effect of genotype, *F*(1,25)=0.0062, p=.94; data not shown). A subset of mice underwent extinction of food responding and cue-induced reinstatement of food seeking. No differences were seen between the groups in the rate of extinction (days to extinction WT: 3.25 ± 0.25; PKMζ KO_constitutive_: 3.38 ± 0.38; t(10)=0.22, p=.83). Both wildtype and PKMζ KO_constitutive_ mice exhibited cue-induced reinstatement of food seeking, however no differences were seen between the groups (effect of session, *F*(1,10)=12.41, p=.006; effect of genotype, *F*(1,10)=0.0007, p=.98; n=6-8/group; Figure 2c).

**Figure 2.**
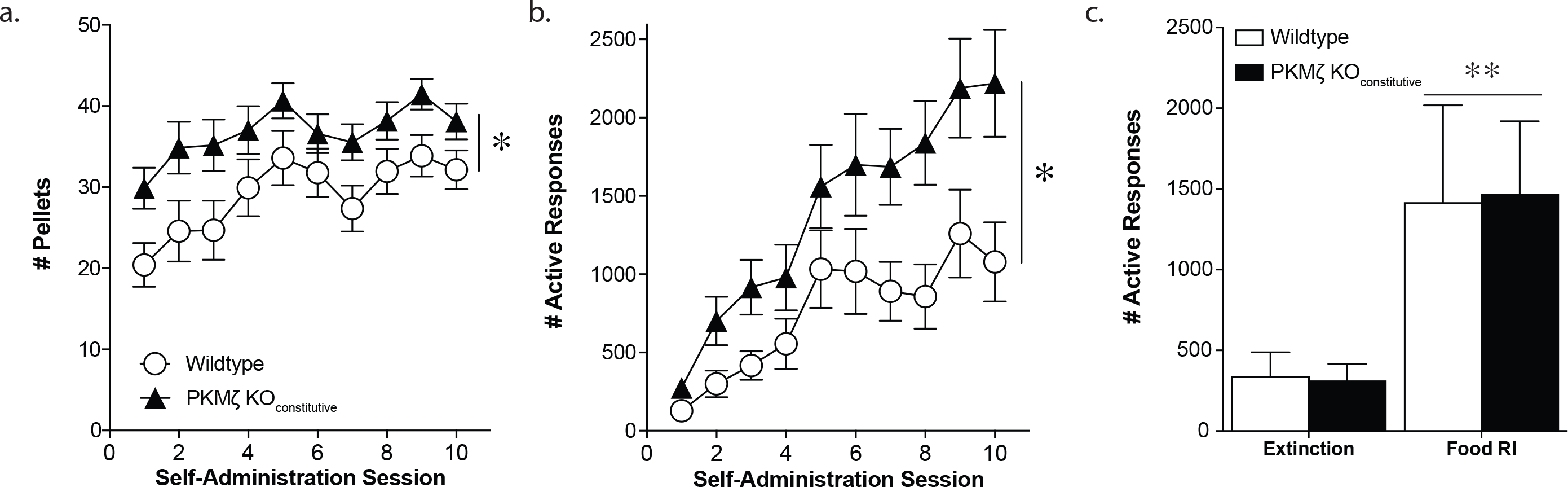
Constitutive PKMζ knockout potentiates food self-administration without altering cue-induced food seeking. PKMζ KO_constitutive_ mice exhibit significantly greater responding (b) and earn more pellets (a) than wildtype mice during a 1 hour food self-administration session. While both wildtype and PKMζ KO_constitutive_ mice exhibit a significant increase in responding during reinstatement (c), there were no differences between the groups. *p<.05, effect of genotype (a,b; n=22/group); **p<.01, effect of session (c; n=6-8/group).

### Constitutive Deletion of PKMζ Potentiates Responding During Cocaine Self-Administration and Cue-Induced Reinstatement of Cocaine Seeking

Following food training, mice received jugular catheterization surgery and underwent ten days of cocaine self-administration. Both wildtype and PKMζ KO_constitutive_ mice showed an increase in cocaine intake over the course of the 10 sessions and no group differences were seen in cocaine intake (effect of session, *F*(9,225)=4.93, p<.0001; effect of genotype, *F*(1,25)=0.41, p=.53; Figure 3a). However, PKMζ KO_constitutive_ mice exhibited higher rates of responding for cocaine compared with wildtype controls (effect of genotype, *F*(1,25)=7.85, p=.009; n=12-15/group; Figure 3b). This effect of genotype was seen in both males and females (Males: effect of genotype, responses= *F*(1,14)=5.43, p=.03; n=7-8/group; Females: effect of genotype, responses= *F*(1,16)=9.38, p=.007; n=9). Following the cocaine self-administration, animals underwent extinction of the cocaine seeking behavior. There was no difference between the two groups in the amount of time taken to reach the extinction criterion (*t*(21)=0.30, p=0.77; Figure 3c). Following extinction, mice were exposed to a cue-induced reinstatement session in which active responses resulted in presentation of the cues previously paired with drug administration in the absence of the drug. Disrupting PKMζ function led to increased cue-induced drug seeking compared with wildtype controls (effect of session, *F*(1,22)=8.71, p=.007; effect of genotype, *F*(1,22)=5.10, p=.03; interaction, *F*(1,22)=4.41, p=.04; Sidak’s multiple comparison, KO vs. WT RI test, adjusted p=.0002; n=8-16/group; Figure 3d).

**Figure 3.**
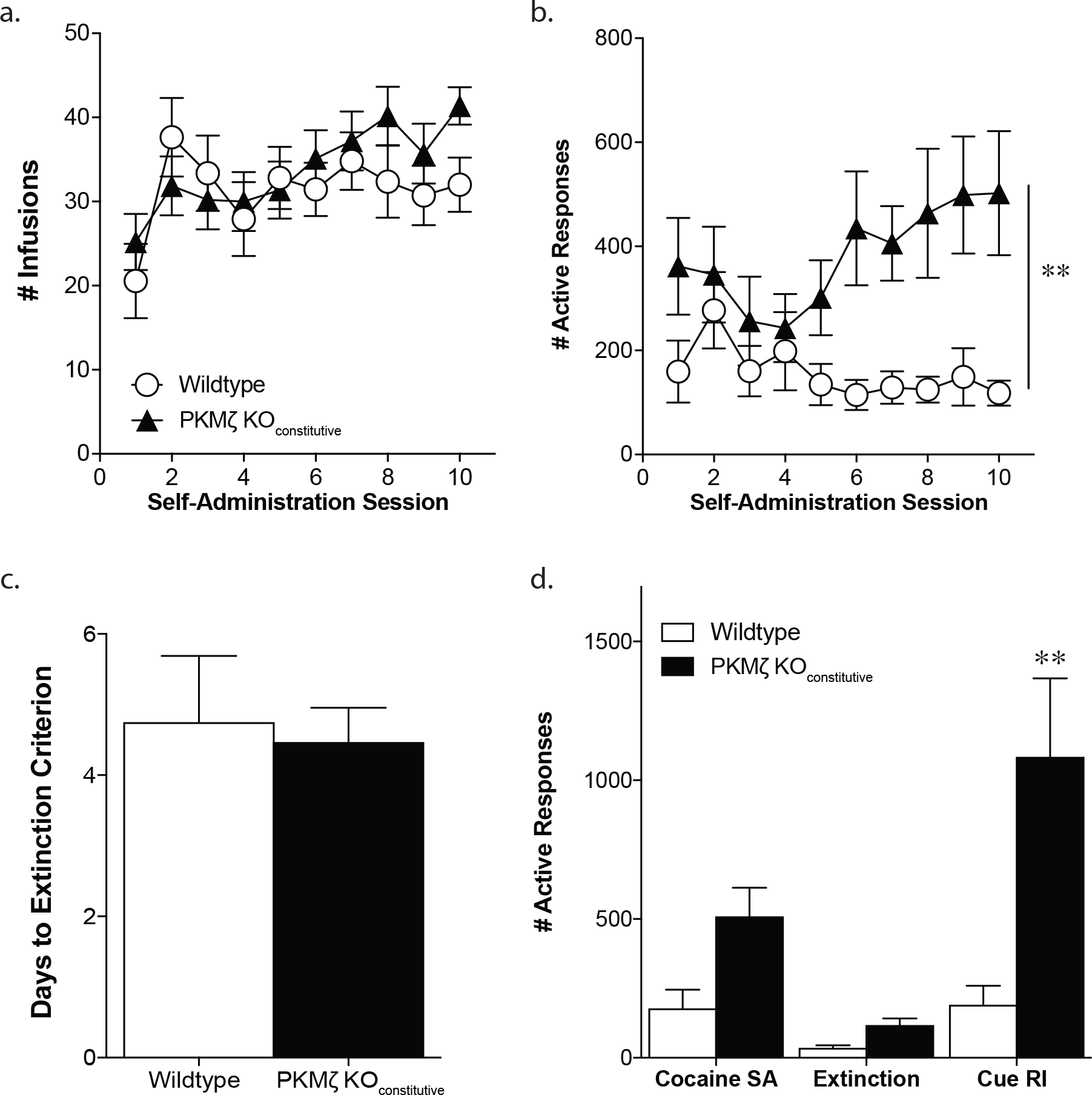
Constitutive PKMζ knockout potentiates cocaine selfadministration and cue-induced cocaine seeking. Although the groups did not disfer in the number of cocaine infusions (a), PKMζ KO_constitutive_ mice exhibited significantly greater responding during both cocaine self-administration (b) and cue-induced reinstatement (d; 2hr session). No differences were seen in the time it took for the groups to extinguish the cocaine seeking behavior (c). **p<.01, effect of genotype (b; n=12-15/group); pairwise comparison WT vs. KO (c; n=8-16/group).

### Site-specific knockdown of PKMζ in the Nucleus Accumbens does not alter Food Seeking or Reinstatement

To determine if the effects of PKMζ deletion on food and cocaine behavior were due to actions of PKMζ within the nucleus accumbens, we injected floxed PKMζ adult mice with an adeno-associated virus expressing either Cre recombinase or GFP into the accumbens. This led to a significant decrease in PKMζ abundance within the nucleus accumbens 6 weeks following viral injection in both males and females [main effect of virus, *F*(1,36)=37.49, p<.0001; Figure 4a]. Of note, the western blot analysis revealed a baseline sex difference with both GFP and Cre injected females exhibiting lower levels of PKMζ abundance compared with males [main effect of sex, *F*(1,36)=7.61, p=.009; Figure 4a]. Similar to what has been shown previously in the PKMζ KO_constitutive_ mice (Volk *et al*, 2013), no changes were seen in the other atypical PKC isozyme, PKCι/λ following PKMζ knockdown [effect of virus, *F*(1,36)=2.05, p=.16; effect of sex, *F*(1,36)=3.48, p=.07; Figure 4b]. Both wildtype and PKMζ KOaccumbens mice showed a gradual increase in the number of pellets earned and active responses per session over the 10 days of food training and no differences were seen between the groups (effect of session, pellets= *F*(9,567)=19.85, p<.0001; responses= *F*(9,567)=26.26, p<.0001; effect of genotype, pellets= *F*(1,63)=0.299, p=.59; responses= *F*(1,63)=0.667, p=.80; N=30-35; Figure 5a,b). Furthermore, no differences were seen between the groups in the rate of extinction (days to extinction WT: 3.5 ± 0.5; PKMζ KO_accumbens_: 4.4 ± 0.62; t(14)=1.007, p=.33). Both wildtype and PKMζ KO_accumbens_ mice exhibited cue-induced reinstatement of food seeking, however no differences were seen between the groups (effect of session, *F*(1, 14)=21.61, p=.0004; effect of genotype, *F*(1,14)=1.24, p=.28; Figure 5c).

**Figure 4.**
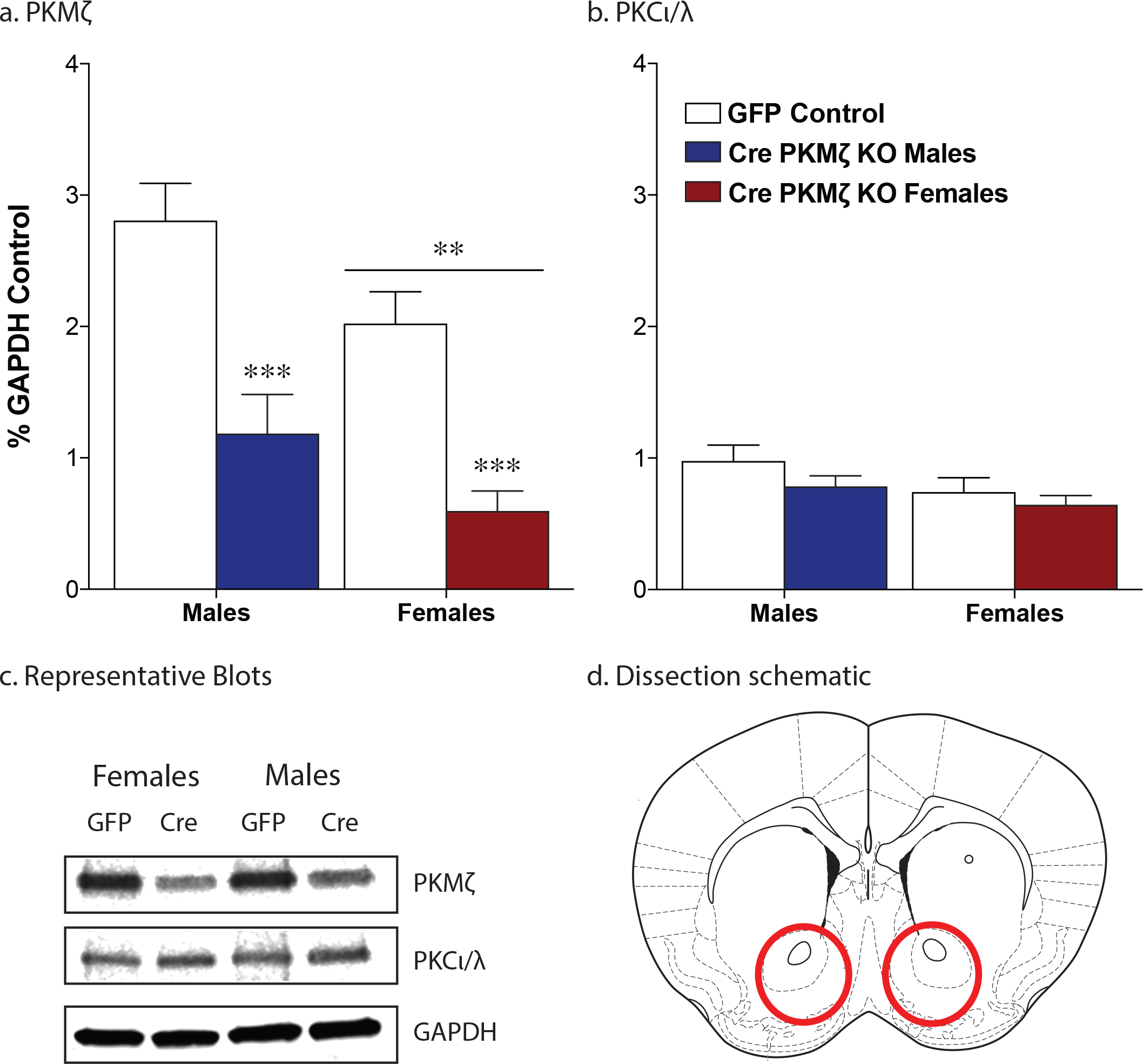
Cre recombinase injection into the nucleus accumbens leads to a significant decrease in PKMζ protein levels without altering PKCι/λ. Quantification of western blot showing a significant decrease in PKMζ protein in the nucleus accumbens following AAV-Cre injection in both male and female mice, as normalized to glyceraldehyde 3-phosphate dehydrogenase (a,c;***p<.0001 main effect of virus; **p<.01 main effect of sex; n=9-13/group). PKMζ knockdown did not alter the levels of PKCi/A within the accumbens (b,c). Red circles indicate area dissected for the confirmation of viral knockdown (d).

**Figure 5.**
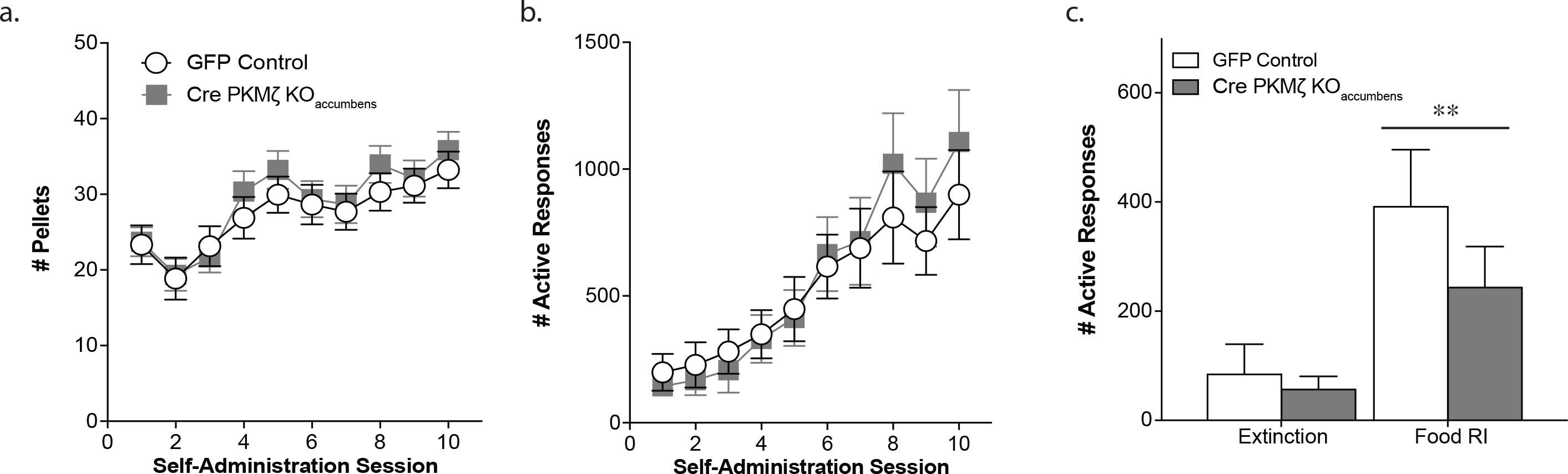
Site-specific knockdown of PKMζ within the nucleus accumbens does not affect food self-administration or reinstatement. No differences were seen between the groups in the number of pellets earned or the number of active responses during the self-administration phase (a, b; n=30-36; 1 hr session). During the cue-induced reinstatement session both groups exhibited a significant increase in responding compared with their extinction responding but no differences were seen between the groups (c; **p<.01 effect of session, n=6-10; 2hr session).

### Site-specific knockdown of PKMζ in the Nucleus Accumbens Potentiates Cocaine Self-Administration in Males but not Females

Male and female wildtype and PKMζ KO_accumbens_ mice showed an increase in cocaine intake over the course of the 10 sessions (males: effect of session, *F*(9,180)=8.99, p<.0001; females: effect of session, *F*(9,243)=13.84, p<.0001; Figure 6a,b). Male PKMζ KO_accumbens_ mice exhibited an increase in cocaine intake and responding for cocaine (effect of genotype, infusions= *F*(1,20)=7.25, p=.01; responses= *F*(1,20)=4.39, p=.049; Figure 6a,c). Furthermore, Male PKMζ KO_accumbens_ mice also exhibited an increase in cue-induced cocaine seeking (effect of session, *F*(1,10)=14.29, p=.0036; effect of viral injection, *F*(1,10)=5.76, p=.037; interaction, *F*(1,10)=4.60, p=.05; Sidak’s multiple comparison, Cre vs. GFP RI test, adjusted p=.0087; n=6-7/group; Figure 6e). However, in the female mice, the local PKMζknockout in the nucleus accumbens had no effect on cocaine self-administration (infusions: *F*(1,27)= 0.489, p=0.490; responses: *F*(1,26)=0.162, p=0.691; n=14-15/group; Figure 6b,c) or reinstatement of cocaine seeking (effect of session, *F*(1,9)=13.39, p=.0052; effect of viral injection, *F*(1,9)=0.13, p=.73; interaction, *F*(1,9)=0.28, p=.61; n=6-8/group; Figure 6f).

**Figure 6.**
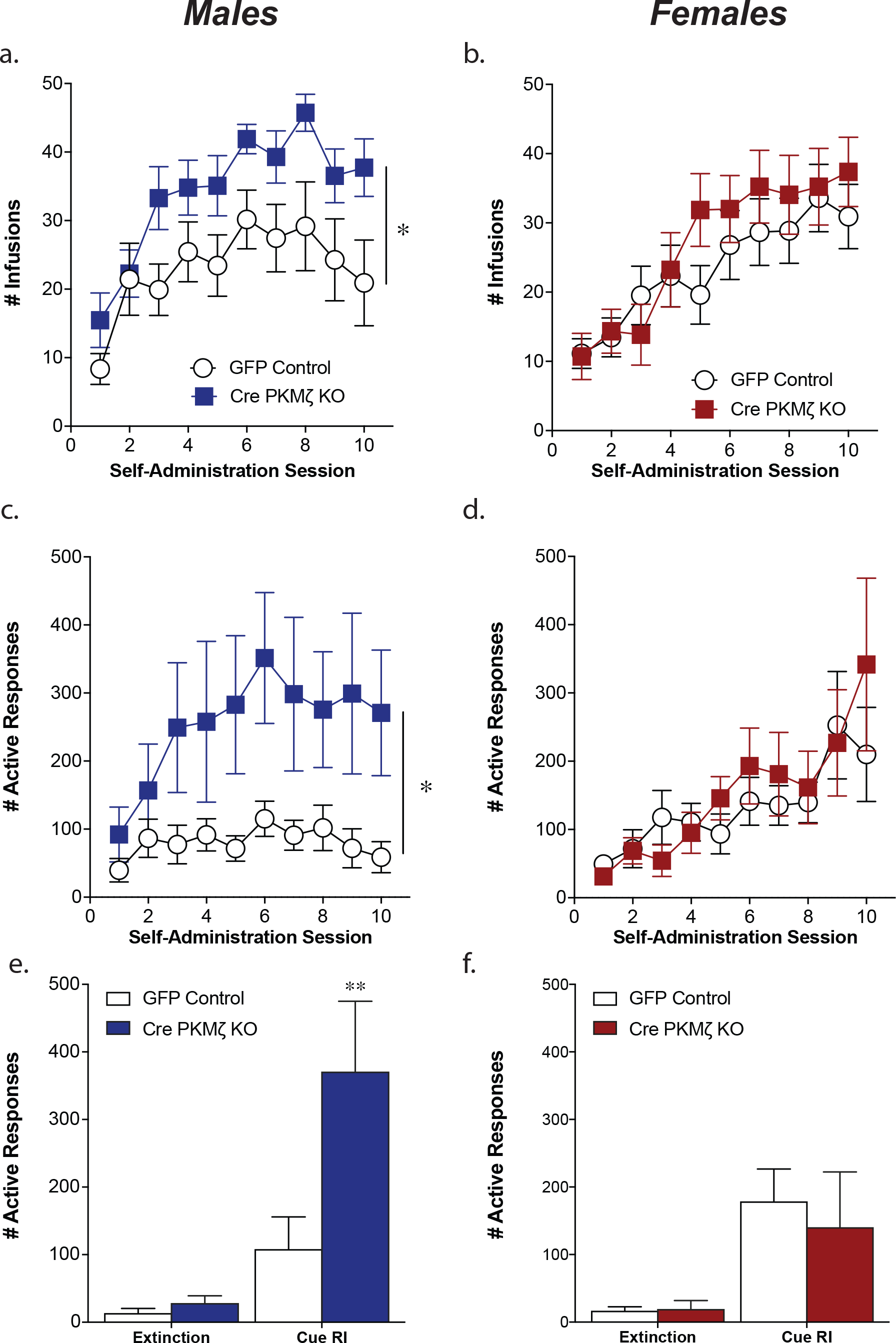
Site-specific knockdown of PKMζ within the nucleus accumbens enhances cocaine self-administration in male mice. Male PKMζ KO_accumbens_ mice exhibited a significant increase in cocaine taking, as indicated by an increase in cocaine infusions earned (a) and active responses completed (c) compared with GFP-injected control males (*p<.05, effect of genotype, n=11/group; 2hr session). Further, male PKMζ KO_accumbens_ mice exhibit greater cocaine seeking during cue-induced reinstatement (e; **p<.01 pairwise comparison GFP vs. Cre on RI day, n=6-7/group). Female PKMζ KO_accumbens_ mice did not differ from GFP injected controls during self-administration (b,d;n=14-15/group) or reinstatement (f, n=6-8/group).

## Discussion

Here, we provide evidence that the atypical PKC, PKMζ, plays a critical role in mediating reward-related behavior. Cocaine self-administration experience led to an increase in PKMζ within the nucleus accumbens at both the transcript and protein levels. This increase appears to be compensatory as we found that constitutive knockout of PKMζ increases cocaine taking and vulnerability to cue-induced cocaine relapse. This increase in cocaine taking and seeking is recapitulated in male mice with a site-specific knockdown of PKMζ in the nucleus accumbens. However, accumbal knockdown of PKMζ did not alter cocaine taking or seeking in females.

### PKMζ Deletion Potentiates Cue-Induced Reinstatement of Cocaine Seeking

The current study is one of two studies to demonstrate a behavioral phenotype in PKMζ knockout mice. Our findings are consistent with previous work demonstrating constitutive PKMζ knockout mice exhibit increased alcohol intake (Lee *et al*, 2014). The data presented here provide support for the idea that PKMζ acts to dampen reward, as cocaine self-administration experience led to an increase in both PKMζ transcript and protein abundance within the nucleus accumbens. This is consistent with increases in PKMζ seen following morphine conditioned place preference (Li *et al*, 2011), a single episode of ethanol drinking (Mulligan *et al*, 2011), and experimenter administered cocaine (Ho *et al*, 2012). However, in the absence of PKMζ mice exhibited an increase in cocaine seeking. This suggests that cocaine-induced increases in PKMζ abundance are compensatory and act to counter the effects of cocaine.

Within the hippocampus, PKMζ enhances AMPA transmission, an effect mediated by NSF-mediated insertion of GluA2-containing AMPA receptors (Yao *et al*, 2008). Additionally, once these GluA2-containing AMPARs are inserted, PKMζ may also make them harder to move out of the synapse (Yu *et al*, 2017). As the removal of GluA2-containing AMPARs and corresponding insertion of GluA2-lacking AMPARs plays a critical role in cocaine craving (Briand *et al*, 2016; Conrad *et al*, 2008; Famous *et al*, 2008; McCutcheon *et al*, 2011), this could provide a mechanism by which PKMζ could dampen reward. As none of the studies examining the relationship between PKMζ and GluA2 were done in the nucleus accumbens, more work is needed to determine whether PKMζ is in fact altering AMPAR subunit composition and how it functions following chronic cocaine exposure.

### The Role of Accumbal PKMζ in Cocaine Seeking is Sex-Specific

The current study was able to recapitulate the increased cocaine seeking seen following constitutive PKMζ knockout with site-specific knockdown of PKMζ within the nucleus accubmens in male mice. However, accumbal knockdown of PKMζ did not alter cocaine taking or seeking in female mice. Of note, wildtype female mice exhibited lower levels of PKMζ abundance compared with males. As these samples were taken after cocaine experience, it is not known whether there are baseline differences in PKMζ abundance or if cocaine alters PKMζ abundance differently in males and females. To date all the studies examining experience-dependent increases in PKMζ abundance have all exclusively examined males (Hsieh *et al*, 2017; King *et al*, 2012; Li *et al*, 2014; Li *et al*, 2011; Xin *et al*, 2014).

There is some evidence that PKMζ has sex-specific effects in other behaviors as well as the cocaine effects reported here. Constitutive deletion of PKMζ leads to decreased anxiety-like behavior in male mice but not females (Lee *et al*, 2013a). However, as anxiolytic medications can decrease cocaine addictive phenotypes (Augier *et al*, 2012; Barbosa-Mendez *et al*, 2017), it is unlikely that our behavioral effects are mediated by this decreased anxiety. PKMζ deletion also increases muscle pain threshold in male mice but not female mice (Nasir *et al*, 2016). Together with the current findings, these studies support a greater role for PKMζ in males. However, as we do see effects of constitutive deletion of PKMζ in females, further work is needed to determine the sex-specific roles of PKMζ in these behaviors.

### Role of PKMζ in food reward

Constitutive PKMζ deletion led to an increase in food self-administration as well along with the increased cocaine taking. This suggests that PKMζ plays a more general role in reward learning and reward consumption. However, systemic administration of a competitive AMPA antagonist decreases sucrose self-administration (Stephens and Brown, 1999) and application of PKMζ in the slice appears to increase AMPAR transmission (Yao *et al*, 2008). In contrast, AMPA antagonist administration into the nucleus accumbens facilitates feeding behavior (Maldonado-Irizarry *et al*, 1995). However, the lack of increase in food seeking following site-specific knockdown of PKMζ suggests that this effect is not accumbally mediated. Thus, PKMζ must play a critical role in food seeking in another brain region. Changes in feeding behaviors have been observed following glutamate manipulations in the ventral tegmental area, the dorsal striatum and the posterodorsal medial amygdala (Alsio *et al*, 2011; D’Souza and Markou, 2011; Rosa *et al*, 2011). Further work will need to be done to determine if the effects seen in the constitutive knockout mice are due to the role of PKMζ in these regions. Despite the increase in food seeking in constitutive knockout mice, they do not demonstrate any alterations in cue-induced reinstatement of food seeking. As many studies manipulating AMPA receptors have seen specific alterations in cocaine seeking without corresponding changes in food seeking (Anderson *et al*, 2008; Briand *et al*, 2016; Briand *et al*, 2014; Famous *et al*, 2008), this supports our hypothesis that PKMζ knockout may be altering accumbal AMPA receptors.

## Conclusion

In the current study, we have shown that constitutive deletion of PKMζ enhances cocaine seeking and taking, while only increasing intake but not seeking of a natural reward. In males, deletion of PKMζ only in the nucleus accumbens is sufficient to enhance cocaine seeking and taking, but has no effect in females. It’s possible that the sex differences observed in this study are also due to sex differences in the glutamatergic system following cocaine exposure.

## Funding and Disclosure

The authors declare no conflict of interest.

This work was supported by National Institute on Drug Abuse (NIDA) Grant R00 DA033372 (L.A.B.) and a Brain & Behavior Research Foundation NARSAD award (L.A.B.).

## Acknowledgements

We thank Dr. Richard L. Huganir for providing the initial breeders for these studies.

